# The sleep gene *insomniac* ubiquitinates targets at postsynaptic densities and is required for retrograde homeostatic signaling

**DOI:** 10.1101/430819

**Authors:** Koto Kikuma, Xiling Li, Sarah Perry, Qiuling Li, Pragya Goel, Catherine Chen, Daniel Kim, Nicholas Stavropoulos, Dion Dickman

**Author notes:** Correspondence: Dion Dickman, Department of Neurobiology, University of Southern California, Los Angeles, CA 90089, USA.

## Abstract

The nervous system confronts challenges during development and experience that can destabilize information processing. To adapt to these perturbations, synapses homeostatically adjust synaptic strength, a process referred to as homeostatic synaptic plasticity. At the *Drosophila* neuromuscular junction, inhibition of postsynaptic glutamate receptors activates retrograde signaling that precisely increases presynaptic neurotransmitter release to restore baseline synaptic strength. However, the nature of the underlying postsynaptic induction process remains enigmatic. Here, we designed a forward genetic screen to identify factors necessary in the postsynaptic compartment to generate retrograde homeostatic signaling. This approach identified *insomniac* (*inc*), a gene that encodes a putative adaptor for the Cullin-3 ubiquitin ligase complex and is essential for normal sleep regulation. Intriguingly, we find that Inc rapidly traffics to postsynaptic densities and is required for increased ubiquitination following acute receptor inhibition. Our study suggests that Inc-dependent ubiquitination, compartmentalized at postsynaptic densities, gates retrograde signaling and provides an intriguing molecular link between the control of sleep behavior and homeostatic plasticity at synapses.

## INTRODUCTION

To maintain stable synaptic activity in the face of stress during development, experience, and disease, the nervous system is endowed with robust forms of adaptive plasticity that homeostatically adjust synaptic strength (Davis and Muller, 2015; Pozo and Goda, 2010; Turrigiano, 2012). The homeostatic control of synaptic plasticity is conserved from invertebrates to humans (Davis, 2013; Frank, 2014; Marder and Goaillard, 2006), and dysfunction in this process is linked to complex neural diseases including Parkinson’s, schizophrenia, Fragile X Syndrome, and autism spectrum disorder (Bourgeron, 2009; Li et al., 2018a; Ramocki and Zoghbi, 2008; Soukup et al., 2018; Wondolowski and Dickman, 2013). Homeostatic adaptations at synapses are expressed through coordinated modulations in the efficacy of presynaptic neurotransmitter release and/or postsynaptic receptor abundance (Burrone and Murthy, 2003; Davis, 2006; Herring and Nicoll, 2016; Malinow, 2003; Perez-Otano and Ehlers, 2005; Shepherd and Huganir, 2007; Turrigiano, 2008; Turrigiano and Nelson, 2004). Although it is apparent that a dialogue involving both anterograde and retrograde trans-synaptic signaling serves to initiate, maintain, and integrate the homeostatic tuning of synaptic strength, the molecular nature of this communication is largely unknown.

The *Drosophila* neuromuscular junction (NMJ) is an established model system to interrogate the genes and mechanisms that mediate the homeostatic stabilization of synaptic strength. At this glutamatergic synapse, genetic loss or pharmacological inhibition of postsynaptic receptors initiates a retrograde signaling system that instructs a compensatory increase in presynaptic neurotransmitter release to restore baseline levels of synaptic strength (Frank et al., 2006; Petersen et al., 1997), a process referred to as presynaptic homeostatic potentiation (PHP). Forward genetic screens in this system have proven to be a powerful tool to identify genes necessary for the expression of PHP (Dickman and Davis, 2009; Frank, 2014; Muller et al., 2011). Work over the past decade has revealed that a rapid increase in both presynaptic Ca^2+^ influx and the size of the readily releasable vesicle pool are necessary to homeostatically enhance neurotransmitter release during PHP (Kiragasi et al., 2017; Li et al., 2018c; Muller and Davis, 2012; Weyhersmuller et al., 2011). Furthermore, candidate molecules involved in retrograde signaling have been proposed (Orr et al., 2017; Wang et al., 2014). However, despite these significant insights, forward genetic screens have failed to shed light on the postsynaptic mechanisms that induce retrograde signaling, a process that remains enigmatic (Chen and Dickman, 2017; Goel et al., 2017; Hauswirth et al., 2018).

Little is known about the signal transduction system in the postsynaptic compartment that initiates retrograde homeostatic communication. It is clear that pharmacological blockade or genetic loss of GluRIIA-containing receptors initiates retrograde PHP signaling. Perturbation of these receptors lead to reduced levels of active (phosphorylated) Ca^2+^/calmodulin-dependent protein kinase II (CaMKII) (Goel et al., 2017; Haghighi et al., 2003; Li et al., 2018c; Newman et al., 2017). However, inhibition of postsynaptic CaMKII activity alone is not sufficient to induce PHP expression (Haghighi et al., 2003), suggesting that additional signaling in the postsynaptic compartment is required to generate retrograde communication. Furthermore, rapid PHP signaling induced by pharmacological receptor blockade does not require new protein synthesis (Frank et al., 2006; Goel et al., 2017). Finally, CaMKII signaling is compartmentalized at postsynaptic densities, where PHP can be expressed with specificity at synapses with diminished receptor function (Li et al., 2018b; Newman et al., 2017), suggesting that retrograde communication happens locally between individual pre- and post-synaptic dyads. Although a role for postsynaptic PI3-cll kinase in PHP was recently proposed (Hauswirth et al., 2018), it is unclear how this signaling is connected to localized glutamate receptor perturbation, compartmentalized changes in CaMKII activity, or retrograde communication to specific presynaptic release sites. Together, these data suggest that translation-independent signaling systems are compartmentalized at postsynaptic densities and function in addition to CaMKII to ultimately drive localized retrograde homeostatic communication to specific presynaptic release sites.

To gain insight into the mechanisms underlying PHP induction, we have designed complementary forward genetic screens to identify genes that specifically function in the postsynaptic compartment to enable retrograde homeostatic signaling. This approach discovered a single gene, *insomniac* (*inc*), that functions in the postsynaptic muscle to induce retrograde communication following postsynaptic glutamate receptor perturbation. *inc* encodes a putative adaptor for the Cullin-3 (Cul3) E3 ubiquitin ligase complex that targets substrates for ubiquitination and is necessary for normal sleep behavior (Pfeiffenberger and Allada, 2012; Stavropoulos and Young, 2011). Our findings suggest that rapid and compartmentalized ubiquitination at postsynaptic densities is a key inductive event during trans-synaptic homeostatic signaling.

## RESULTS

### Electrophysiology-based forward genetic screens identify *inc*

We first generated a list of ~800 neural and synaptic genes to screen for defects in the ability to express PHP. A substrantial portion of these genes were gleaned from various studies linking genes to schizophrenia, intellectual disability, autism, and Fragile X Syndrome (see Methods for more details). We hypothesized that these genes and targets of the Fragile X Mental Retardation Protein (FMRP) might provide a rich source to assess for postsynaptic roles in homeostatic synaptic signaling. First, previous studies have established intriguing links between homeostatic plasticity and complex neurological and neuropsychiatric diseases (Ramocki and Zoghbi, 2008; Wondolowski and Dickman, 2013). Second, FMRP itself has important roles at postsynaptic densities (Muddashetty et al., 2007; Schutt et al., 2009; Tsai et al., 2012) and has been implicated in homeostatic signaling at mammalian synapses (Henry, 2011; Lee et al., 2018; Soden and Chen, 2010; Zhang et al., 2018). We established a list of *Drosophila* genes that are homologs of the ~800 genes from the initial gene list (see Methods and Supplementary Table 1), and obtained 134 genetic mutants and 284 RNAi lines in *Drosophila* representing these genes to screen for potential defects in PHP expression (Fig. 1a, d).

We used two distinct ways to screen mutants and RNAi lines for their effects on PHP expression. To screen the 134 mutants, we used an established screening approach that utilizes a rapid pharmacological assay to assess PHP (Dickman and Davis, 2009; Muller et al., 2011). In this assay, application of the postsynaptic glutamate receptor antagonist philanthotoxin (PhTx) inhibits miniature neurotransmission, but synaptic strength (evoked EPSPs) remains similar to baseline values because of a homeostatic increase in presynaptic neurotransmitter release (quantal content). For each mutant, we quantified synaptic strength following 10 min incubation in PhTx (Fig. 1b). This led to the identification of twelve potential PHP mutants with significantly reduced synaptic strength after PhTx application, indicative of either reduced baseline transmission or a failure to express PHP. Next, baseline transmission was assessed in these mutants by recording in the absence of PhTx; six mutants with reduced baseline neurotransmission were identified and not studied further (Supplementary Table 2). The remaining six mutants represent genes necessary to express PHP (Fig. 1a, b), including the active zone component *fife*, which was recently shown to be necessary for PHP expression (Bruckner et al., 2017).

**Figure 1:**
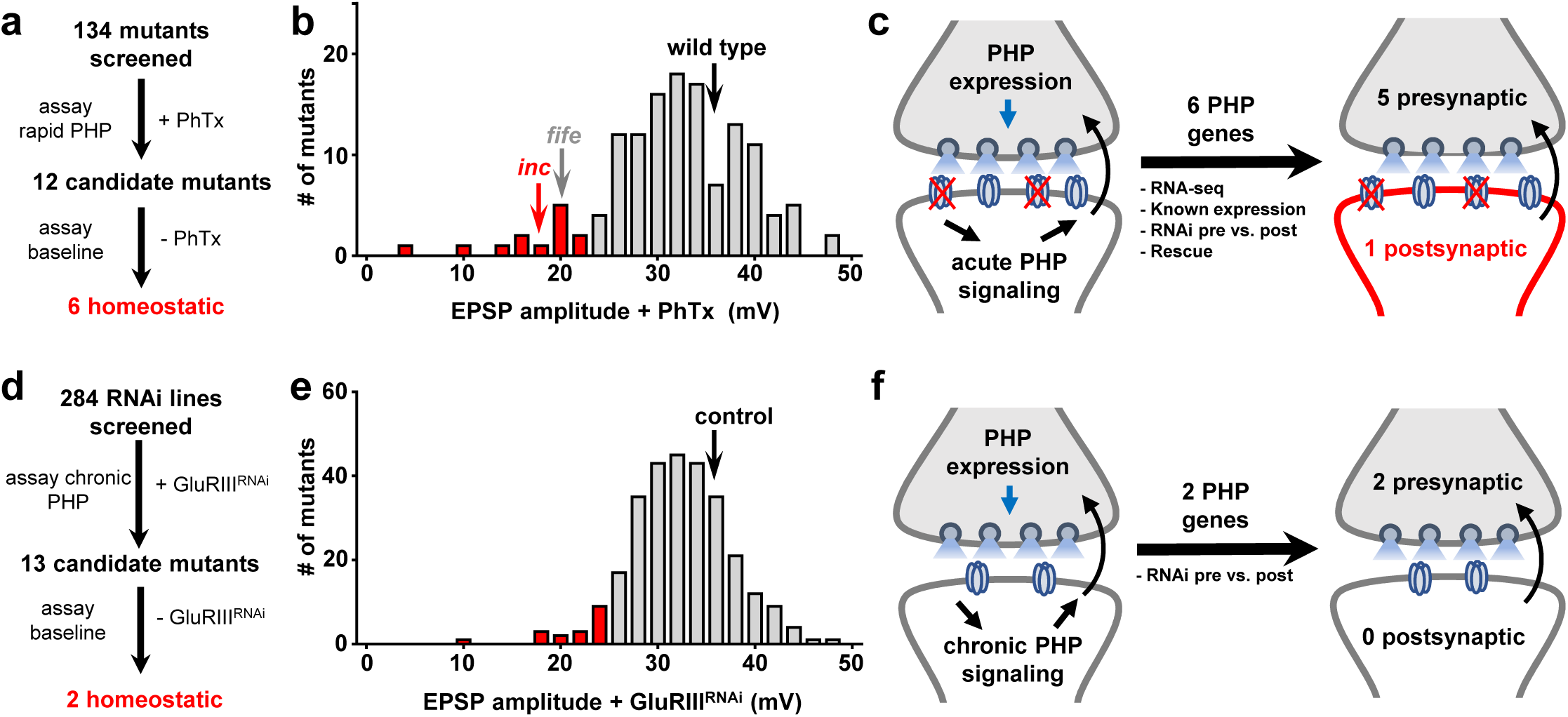
Dual forward genetic screens identify 6 genes necessary for PHP induction and/or expression in distinct synaptic compartments. (**a** and **d**) Electrophysiology-based forward genetic screening strategy and outcomes for the PhTx (a) and *GluRIII* knock down (d) approaches. (**b** and **e**) Average EPSP amplitudes of each mutant or RNAi line screened following PhTx application (b) or *GluRIII* knock down (e). In wild-type controls, inhibition of glutamate receptors results in reduced mEPSP amplitude, as expected. However, EPSP amplitude remains similar to baseline values due to a homeostatic increase in presynaptic neurotransmitter release (quantal content). Highlighted in red are all mutants that showed EPSP values > two standard deviations below controls. (**c** and **f**) Determination of pre- and post-synaptic functions for the positive hits from the screens.

In parallel, we assessed PHP in the 284 RNAi lines using an established stock that drives the RNAi transgene in both neurons and muscle. Importantly, postsynaptic glutamate receptor expression is also reduced in this stock through RNAi-mediated knock-down of the GluRIII receptor transcript (Brusich et al., 2015). After crossing each RNAi line to this stock, we quantified electrophysiological recordings and identified 13 genes that were putatively necessary for PHP expression (Fig. 1d). To determine baseline synaptic strength in these RNAi lines, we expressed each in neurons and muscle in the absence of GluRIII knock down. Of these thirteen genes, eleven exhibited a significant decrease in EPSP amplitude after crossing to the control stock, suggesting reduced baseline transmission (Supplementary Table 2). In contrast, two RNAi lines displayed normal baseline synaptic strength, indicating they were specifically necessary for PHP expression (Fig. 1d, e). Importantly, these two genes targeted by RNAi lines were also identified in the PhTx screen, validating this complementary screening strategy. Together, these two screens identified six genes whose requirement for PHP has not been previously described.

If a gene functioned in the presynaptic neuron, this would imply that it was involved in the expression of increased neurotransmitter release characteristic of PHP, while a postsynaptic function in the muscle would suggest an involvement in the induction of PHP signaling. We therefore used several strategies to determine in which synaptic compartment each gene was required for the induction or expression of PHP. For each of these six genes, we assessed RNA-seq expression profiles (Chen and Dickman, 2017), known expression patterns, genetic rescue and/or tissue-specific RNAi knockdown (Fig. 1c, f). Together, this analysis revealed five genes that function in the presynaptic neuron, and only a single gene, *insomniac* (*inc*), that functions in the postsynaptic cell (Fig. 1c, f). Given that the postsynaptic mechanisms that drive the induction of PHP are poorly defined, we focused on characterizing the role of *inc* in PHP signaling.

### *inc* is required in the postsynaptic muscle to drive retrograde PHP signaling

To further investigate the role of *inc* in PHP, we first generated new null alleles using CRISPR/Cas9 genome editing technology (Gratz et al., 2013a). We obtained two independent mutations in the *inc* locus causing premature stop codons (Fig. 2a), alleles we named *inc^kk3^* and *inc^kk4^*. We confirmed that both alleles are protein nulls by immunoblot analysis with an anti-Inc antibody (Fig. 2b). Furthermore, behavioral analysis demonstrated that both *inc^kk^* mutants exhibit severely shortened sleep, similar to previously described *inc* null alleles (Supplementary Fig. 1 and (Stavropoulos and Young, 2011)).

**Figure 2:**
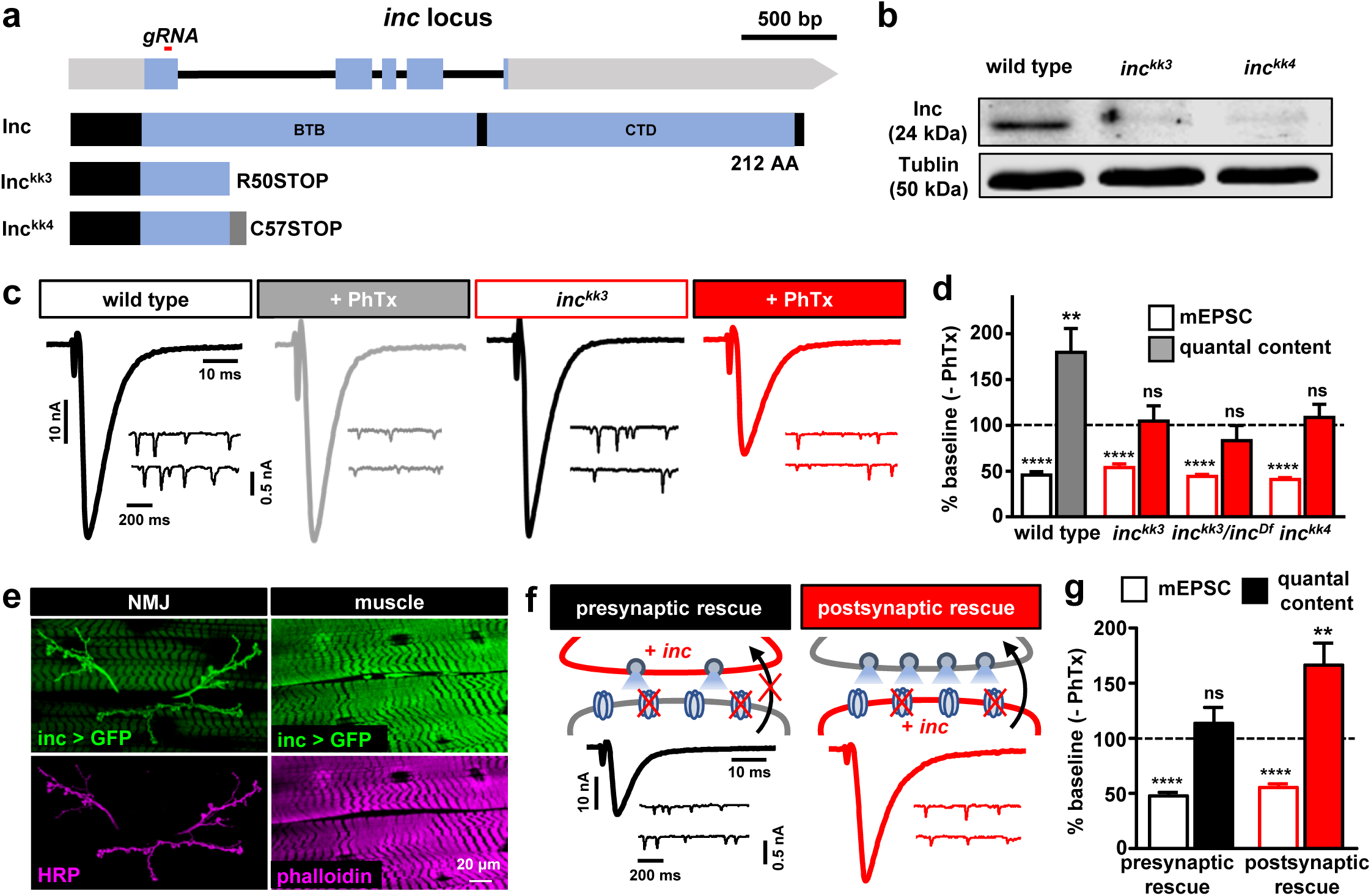
*inc* is required in the postsynaptic muscle to drive retrograde PHP signaling. **(a)** Schematic of the *Drosophila inc* locus, with the region targeted by the guide RNA to generate *inc^kk3^* and *inc^kk4^* shown. (Bottom) Structure of Inc and the predicted Inc mutant proteins. **(b)** Anti-Inc immunoblot analysis from whole adult lysates confirms that both *inc^kk3^* and *inc^kk4^* are protein null alleles. **(c)** Acute expression of PHP requires *inc*. Representative EPSC and mEPSC traces for wild type (*w^1118^*) and *inc^kk3^* mutants before and after PhTx application. *inc^kk3^* mutants fail to homeostatically increase presynaptic neurotransmitter release after PhTx application, resulting in reduced EPSC amplitudes. **(d)** Average mEPSC amplitude and quantal content values following PhTx application relative to baseline (-PhTx) are shown for the indicated genotypes. **(e)** Representative NMJ images of GFP expression driven by the *inc* promoter (*inc-Gal4;UAS-eGFP/+*). Anti-HRP (neuronal membrane marker) and anti-phalloidin (muscle actin marker) are shown. Inc is expressed in both presynaptic neurons and postsynaptic muscles. **(f)** Schematic and representative EPSC and mEPSC traces in which *UAS-smFP-inc* is expressed in motor neurons (presynaptic rescue: *inc^kk3^;Ok371-Gal4/UAS-smFP-inc*) or muscle (postsynaptic rescue: *inc^kk3^; UAS-smFP-inc/+; MHC-Gal4/+*) in *inc* mutant backgrounds following PhTx application. Postsynaptic expression of *inc* fully restores PHP expression, while PHP fails in the presynaptic rescue condition. **(g)** Average mEPSC and quantal content values in (f) relative to baseline. Error bars indicate ±SEM.

Next, we characterized synaptic physiology in *inc* mutants using two-electrode voltage clamp recordings. We first confirmed that baseline synaptic transmission was largely unperturbed by the loss of *inc* (Fig. 2c and Supplementary Fig. 1). However, while PhTx application reduced mEPSC amplitudes in both wild type and *inc* mutants, no homeostatic increase in presynaptic release was observed in *inc* mutants, resulting in reduced EPSC amplitude (Fig. 2c, d). Similar results were found for *inc^kk3^/inc^Df^* and *inc^kk4^* mutants (Fig. 2d and Supplementary Table 3). In addition, *inc* mutants failed to express PHP over chronic time scales when combined with *GluRIIA* mutations (Supplementary Fig. 2). Thus, *inc* is necessary for the expression of PHP over both acute and chronic time scales.

If *inc* were required in the neuron for PHP expression, this would indicate a function in augmenting presynaptic neurotransmitter release. In contrast, if *inc* were required in the muscle, this would suggest a role in postsynaptic retrograde communication. We therefore determined in which synaptic compartment *inc* is required for PHP expression. First, we used an *inc-Gal4* transgene (Stavropoulos and Young, 2011) to express a GFP reporter and observed the GFP signal in both presynaptic motor neurons and the postsynaptic musculature (Fig. 2e), as previously described (Li et al., 2017). To determine in which compartment *inc* expression was required for PHP, we performed a tissue-specific rescue experiment using a UAS transgene expressing Inc fused to a *spaghetti monster* Fluorescent Protein (smFP) 10xFlag tag ((*UAS-smFP-inc;* (Viswanathan et al., 2015)). Consistent with the notion that smFP-Inc does not antagonize endogenous Inc, overexpression of this transgene had no impact on baseline synaptic transmission or PHP expression (Supplementary Fig. 3). Expression of this transgene with *inc-Gal4* also rescued the sleep deficits in *inc* mutants (Supplementary Fig. 1), suggesting that *smFP-inc* recapitulates Inc function. Importantly, PHP expression was fully restored in *inc* mutants when this transgene was expressed specifically in the postsynaptic muscle, but not when expressed in the presynaptic neuron (Fig. 2f, g). These experiments indicate that *inc* function in the postsynaptic muscle is sufficient to enable retrograde PHP signaling.

### Inc functions downstream of CaMKII or in a parallel pathway to generate retrograde PHP signaling

To characterize the postsynaptic functions of *inc* that enable PHP signaling, we sought to define at what point *inc* is required in this process. First, we assessed whether *inc* mutants exhibit alterations in the localization or abundance of postsynaptic glutamate receptors, key components that initiate PHP signaling. However, we found no significant difference in glutamate receptor puncta signal intensity or localization (Fig. 3a, b).

**Figure 3:**
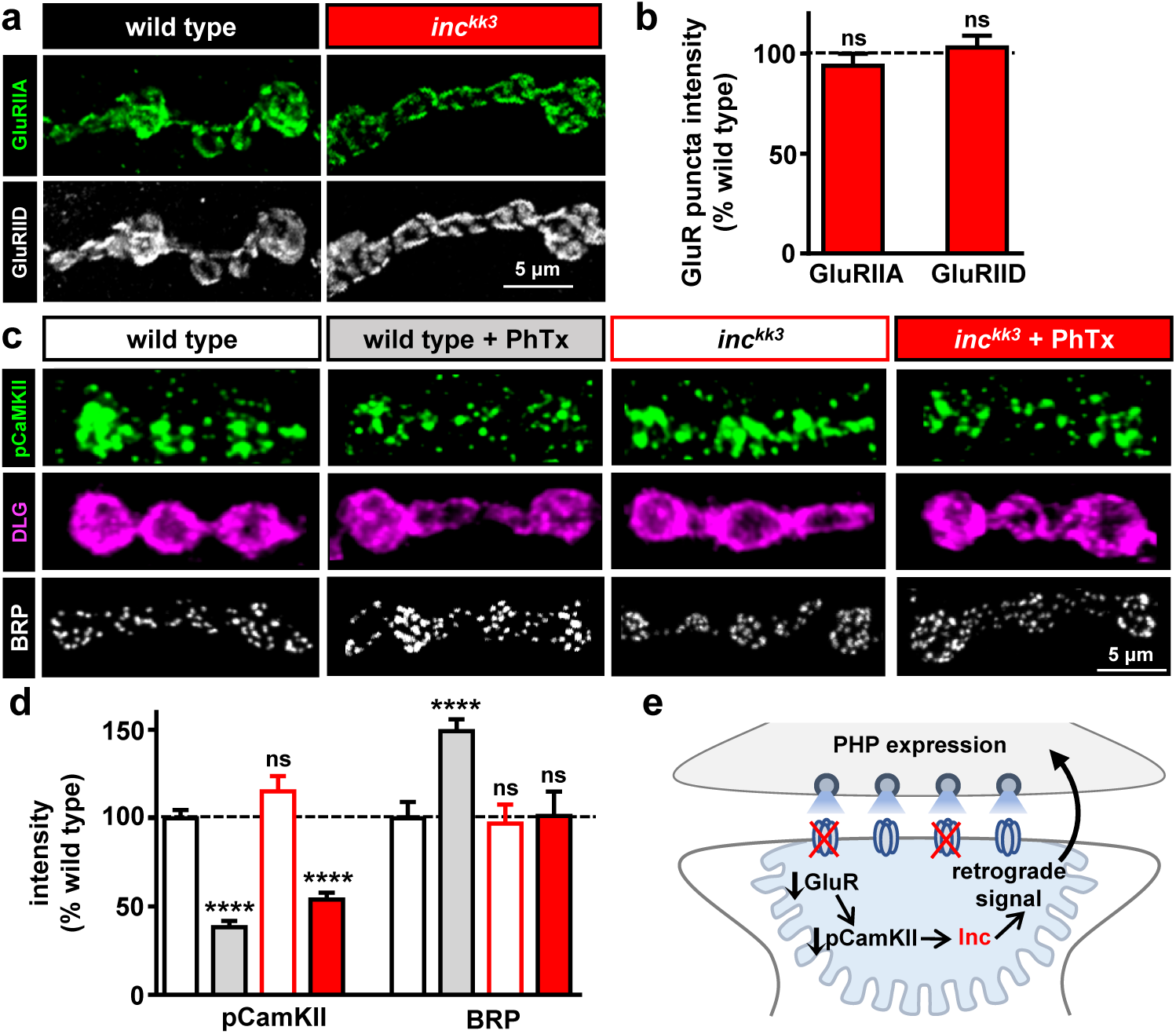
*inc* functions downstream of or in parallel to CaMKII but upstream of retrograde PHP signaling. **(a)** Representative images from wild type and *inc^kk3^* NMJs immunstained with antibodies against the postsynaptic glutamate receptor subunits GluRIIA and GluRIID. No alteration in glutamate receptor levels is observed in *inc* mutants. **(b)** Quantification of average intensity levels of GluRIIA and GluRIID. **(c)** Representative NMJ images of wild type and *inc* mutants immunostained with anti-pCaMKII, -DLG (Discs Large) and -BRP before and after PhTx application. A similar reduction in pCaMKII levels are observed following PhTx application in both wild type and *inc* mutants. In contrast, BRP levels are increased after PhTx application in wild type, but do not change after PhTx application to *inc* mutants, consistent with a lack of presynaptic PHP expression. **(d)** Quantification of average intensity levels of pCaMKII and BRP after PhTx application relative to wild-type values in the indicated genotypes. **(e)** Schematic illustrating *inc* involvement in retrograde PHP signaling pathways. Error bars indicate ±SEM.

Next, we examined the compartmentalized reduction in CaMKII activity, thought to be a key inductive event during retrograde PHP signaling. Indeed, postsynaptic expression of a constitutively active form of CaMKII occludes PHP expression (Haghighi et al., 2003; Li et al., 2018b), and reduced pCaMKII immunofluorescence intensity at the postsynaptic density is observed following loss or pharmacological blockade of glutamate receptors (Goel et al., 2017; Li et al., 2018b; Newman et al., 2017). Further, because the ubiquitination of Inc substrates could trigger their proteolysis, we considered whether Inc might degrade CaMKII following glutamate receptor perturbation. If so, *inc* mutants might fail to reduce pCaMKII abundance at postsynaptic densities following PhTx application, a process thought to be necessary to enable retrograde PHP signaling (Haghighi et al., 2003; Li et al., 2018b). However, pCaMKII levels were similar at the NMJs of *inc* mutants and wild type controls in baseline conditions, and were also reduced to similar levels following PhTx application (Fig. 3c, d).

Finally, retrograde signaling from the postsynaptic compartment following PHP induction leads to remodeling of the presynaptic active zone scaffold bruchpilot ((BRP; (Goel et al., 2017; Weyhersmuller et al., 2011)). We therefore determined whether BRP is remodeled in *inc* mutants following PhTx application. As expected, BRP puncta intensity rapidly increased at presynaptic terminals following PhTx application at wild-type NMJs. However, no change in BRP puncta levels was observed in *inc* mutants following PhTx application (Fig. 3c, d). Together, these results demonstrate that *inc* functions downstream of or in parallel to CaMKII in the postsynaptic compartment, where it is necessary for the retrograde homeostatic signaling that adaptively modulates presynaptic structure and neurotransmitter release (schematized in Fig. 3e).

### Endogenously tagged Inc is rapidly transported to postsynaptic densities following glutamate receptor perturbation

*inc* encodes a highly conserved protein with homology to the Bric-à-brac, Tramtrack, and Broad/Pox virus zinc finger (BTB/POZ) superfamily, which includes adaptors for the Cul3 E3 ubiquitin ligase complex. Inc physically interacts with Cul3, and Cul3 is similarly required for normal sleep, implicating Inc as a substrate adaptor for the Cul3 complex (Pfeiffenberger and Allada, 2012; Stavropoulos and Young, 2011). Consistent with such a mechanism, we observed that knock-down of *Cul3* in muscle, but not in neurons, disrupted the expression of PHP after PhTx application (Supplementary Fig. 4), suggesting that Cul3 works with Inc in the postsynaptic compartment to drive retrograde PHP signaling. The Cul3-Inc complex might ubiquitinate substrates and cause their degradation by the proteasome. Alternatively, Cul3-Inc may regulate substrates by non-degradative mechanisms, including mono-ubiquitination (schematized in Fig. 4b), a post-translational modification that can modulate protein trafficking and signaling (Jin et al., 2012; Kobayashi et al., 2004; Lu and Pfeffer, 2014). A recent study rigorously explored the role of proteasomal degradation during PHP at the *Drosophila* NMJ (Wentzel et al., 2018). Postsynaptic PHP signaling was not impacted by acute pharmacological or chronic genetic inhibition of proteasome-mediated protein degradation (Wentzel et al., 2018). These data and our findings therefore suggest that Cul3 and Inc may mono-ubiquitinate substrates in the postsynaptic compartment to trigger rapid PHP signaling.

**Figure 4:**
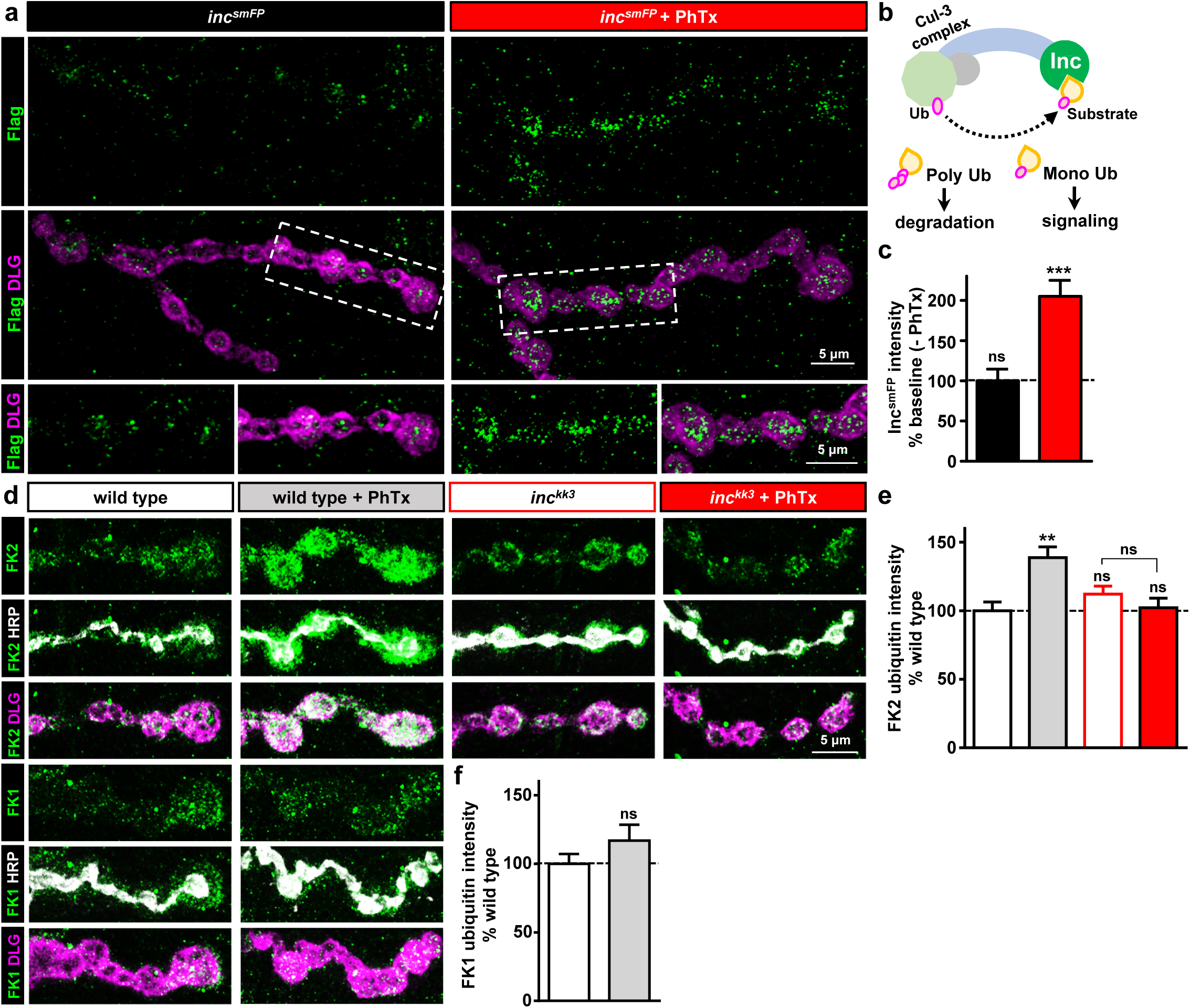
Endogenously tagged Inc rapidly traffics to postsynaptic densities and promotes ubiquitination following glutamate receptor perturbation. **(a)** Representative NMJ images of endogenously tagged Inc^smFP^ before and after PhTx application. NMJs are immunostained with anti-FLAG and the postsynaptic scaffold marker DLG. Areas outlined by dashed-line are shown at higher magnification below. **(b)** Schematic of the Cul3-Inc ubiquitin ligase complex that targets substrates for mono- and poly-ubiquitination. **(c)** Quantification of Inc^smFP^ intensity within the postsynaptic density (marked by DLG) following PhTx application relative to baseline (-PhTx). **(d)** Representative NMJ images from wild type and *inc* mutants immunostained with anti-FK2 (mono- and poly-Ubiquitin), anti-FK1 (poly-Ubiquitin only), the postsynaptic density marker DLG, and the presynaptic membrane marker HRP, before and after PhTx application. At wild-type NMJs, the FK2 signal rapidly increases at postsynaptic densities (indicated by the signal outside of HRP) after PhTx application, while no change is observed in the FK1 signal. However, no change in FK2 intensity is observed at baseline or after PhTx application of *inc* mutant NMJs. Quantification of average FK2 **(e)** and FK1 **(f)** immunointensity after PhTx application, normalized to wild type levels. Error bars indicate ±SEM.

A localized reduction in active CaMKII is observed specifically at the postsynaptic density following genetic loss or pharmacological perturbation of glutamate receptors ((Goel et al., 2017; Newman et al., 2017) and Fig. 3)), suggesting that the key processes driving synapse-specific retrograde PHP signaling occur in this structure(Li et al., 2018b). We therefore first determined whether Inc is present at the postsynaptic density. We endogenously tagged *inc* with an smFP tag (*inc^smFP^;* see Methods) and verified that this tag does not disrupt basal synaptic transmission or PHP expression (Supplementary Fig. 1 and 3). Imaging of Inc^smFP^ at the larval NMJ revealed a low and diffuse cytosolic signal with some enrichment at postsynaptic densities (Fig. 4a). Strikingly, we found that the intensity of Inc^smFP^ was rapidly enhanced at postsynaptic densities after perturbation of glutamate receptors using 10 min application of PhTx (Fig. 4a, c).

Next, we immunostained the NMJ with two anti-Ubiquitin antibodies at basal conditions and following 10 min PhTx incubation in wild type and *inc* mutants. The FK2 antibody recognizes both poly- and mono-ubiquitinylated proteins (Fujimuro et al., 1994; Wentzel et al., 2018), while the FK1 antibody recognizes only poly-ubiquitinylated conjugates (Fujimuro et al., 1994; Ma et al., 2016). We found that the ubiquitin signal labeled by FK2 rapidly increased at postsynaptic densities following PhTx application, while no change in the FK1 signal was observed (Fig. 4d, e). This suggests that acute glutamate receptor perturbation increases mono-ubiquitination at postsynaptic densities. However, no change in the FK2 signal was observed at postsynaptic densities in *inc* mutants following PhTx application (Fig. 4d, e), indicating that *inc* is required for the rapid and compartmentalized increase in ubiquitinated substrates following postsynaptic glutamate receptor perturbation. Thus, Inc rapidly traffics to postsynaptic densities and may locally target substrates for ubiquintation within minutes of glutamate receptor blockade during retrograde homeostatic signaling.

## DISCUSSION

By screening more than 300 synaptic genes, we have identified *inc* as a key postsynaptic regulator of retrograde homeostatic signaling at the *Drosophila* NMJ. Our data suggest that Inc is recruited to the postsynaptic density within minutes of glutamate receptor perturbation, where it promotes local ubiquitination. *inc* functions downstream of or in parallel to CaMKII and upstream of retrograde signaling during PHP. Together, our findings implicate a post-translational signaling system involving mono-ubiquitination in the induction of retrograde homeostatic signaling at postsynaptic compartments.

Although forward genetic screens have been very successful in identifying genes required in the presynaptic neuron for the expression of PHP, these screens have provided less insight into the postsynaptic mechanisms that induce retrograde homeostatic signaling. It seems clear that many genes acting presynaptically are individually required for PHP expression (Bruckner et al., 2017; Dickman and Davis, 2009; Dickman et al., 2012; Kiragasi et al., 2017; Muller et al., 2012; Muller et al., 2011; Tsurudome et al., 2010; Younger et al., 2013), with loss of any one completely blocking PHP expression. In contrast, forward genetic screens have largely failed to uncover new genes functioning in the postsynaptic muscle during PHP, implying some level of redundancy. The specific postsynaptic induction mechanisms driving retrograde PHP signaling have therefore remained unclear (Chen and Dickman, 2017; Goel et al., 2017), further complicated by cap-dependent translation and metabolic pathways somehow engaging with postsynaptic PHP signaling over chronic, but not acute, time scales (Kauwe et al., 2016; Penney et al., 2012; Penney et al., 2016). Therefore, it is perhaps not surprising that despite screening hundreds of mutants, we found only a single gene, *insomniac*, to be required for PHP induction. Inc is expressed in the nervous system and can traffic to the presynaptic terminals of motor neurons (Li et al., 2017). In the context of PHP signaling, however, we found *inc* to be required in the postsynaptic compartment, where it functions downstream of or in parallel to CaMKII. One attractive possibility is that a reduction in CaMKII-dependent phosphorylation of postsynaptic targets enables a subsequent ubiquitination by Cul3-Inc complexes, and that this modification ultimately drives retrograde signaling during PHP. Indeed, reciprocal influences of phosphorylation and ubiquitination on common targets are a common regulatory feature in a variety of signaling systems (Haglund and Dikic, 2005; Karin and Ben-Neriah, 2000; Kawabe and Brose, 2011). This dynamic interplay of phosphorylation and ubiquitination in the postsynaptic compartment may enable a sensitive and tunable mechanism for controlling the timing and calibrating the amplitude of retrograde signaling at the NMJ.

The substrates targeted by Inc for ubiquitination during PHP induction are not known, but there are some candidate pathways to assess. One possibility is that Inc could regulate membrane trafficking events important for retrograde signaling. Multiplexin, a fly homolog of collagen XV/XVIII, and Semaphorin 2B, a secreted protein, were recently proposed to function in retrograde PHP signaling (Orr et al., 2017; Wang et al., 2014), as was postsynaptic endosomal trafficking through Rab11 (Hauswirth et al., 2018). While the relationship between these factors and whether and to what extent trafficking and secretion are regulated during PHP signaling are unclear, the functions of these proteins could be modulated by Cul3- and Inc-dependent ubiquitination. Interestingly, Cul3-dependent mono-ubiquitination regulates membrane trafficking in a Ca^2^+-dependent manner (Jin et al., 2012; McGourty et al., 2016), and Inc could plausibly modulate membrane trafficking during retrograde signal transduction. Alternatively, postsynaptic scaffolds and glutamate receptors may be key Inc substrates at the *Drosophila* NMJ, given that these proteins are targets for ubiquitin-mediated signaling and remodeling at mammalian dendritic spines (Burbea et al., 2002; Colledge et al., 2003; Foot et al., 2017; Hicke and Dunn, 2003; Lin and Man, 2013; Schwarz et al., 2010). Indeed, there is evidence that signaling complexes composed of neurotransmitter receptors, CaMKII, and membrane-associated guanylate kinases are intimately associated at postsynaptic densities in *Drosophila* (Gillespie and Hodge, 2013; Hodge et al., 2006; Lu et al., 2003), and there has been speculation that these complexes are targets for modulation during PHP signaling (Goel et al., 2017; Newman et al., 2017). Ubiquitin-mediated signaling is therefore an attractive process to mediate the translation-independent induction PHP, given the rapid and local enrichment of ubiquitinated proteins triggered at postsynaptic densities following glutamate receptor perturbation and the variety of potential targets sequestered in these compartments.

Although it is well established that the ubiquitin proteasome system can sculpt and remodel synaptic architecture, the importance of mono-ubiquitination at synapses is less well studied. Ubiquitin-dependent pathways play key roles in synaptic structure, function, and degeneration, while also contributing to activity-dependent dendritic growth (DiAntonio et al., 2001; Ehlers, 2003; Hamilton et al., 2012; Hamilton and Zito, 2013; Tai and Schuman, 2008; Tian and Wu, 2013; Wan et al., 2000; Wang et al., 2017). However, the fact that some proteins persist for long periods at synapses suggests that any modification of these proteins by ubiquitin might be non-degredative and reversible. Indeed, a recent study revealed a remarkable heterogeneity in the stability of synaptic proteins, with some short lived and rapidly turned over, while others persist for long times scales and are extremely stable, with half lives of months or longer (Heo et al., 2018). At the *Drosophila* NMJ, rapid ubiquitin-dependent proteasomal degradation at presynaptic terminals is necessary for the expression of PHP through modulation of the synaptic vesicle pool (Wentzel et al., 2018). In contrast, postsynaptic proteasomal degradation is not required for rapid PHP signaling, suggesting that ubiquitin-dependent pathways in the postsynaptic compartment contribute to PHP signaling by non-degradative mechanisms. Our data demonstrate that Cul3 and Inc function in muscle is necessary to enable retrograde PHP signaling, and suggest that these proteins trigger rapid mono-ubiquitination at postsynaptic densities. Interestingly, synaptic proteins can be ubiquitinated in less than 15 seconds following depolarization-induced Ca^2^+ influx at synapses (Chen et al., 2003). Together, both poly- and mono-ubiquination may function in combination with other rapid and reversible processes, including phosphorylation, at pre- and post-synaptic compartments to enable robust and diverse signaling outcomes during plasticity.

A prominent hypothesis postulates that a major function of sleep is to homeostatically regulate synaptic strength following experience-dependent changes that accrue during wakefulness (Tononi and Cirelli, 2014; Vyazovskiy and Harris, 2013). Several studies have revealed provocative changes in neuronal firing rate and synaptic structure and during sleep/wake behavior (Bushey et al., 2011; de Vivo et al., 2017; Diering et al., 2017; Gilestro et al., 2009; Hengen et al., 2016; Li et al., 2017; Yang et al., 2014), yet few molecular mechanisms linking the electrophysiological process of homeostatic synaptic plasticity and sleep have been identified. Our finding that *inc* is required for the homeostatic control of synaptic strength provides an intriguing link to earlier studies which implicated *inc* in the regulation of sleep (Pfeiffenberger and Allada, 2012; Stavropoulos and Young, 2011). It remains to be determined whether the role of *inc* in controlling PHP signaling at the NMJ is related to the impact of *inc* on sleep and whether Inc targets the same substrates to regulate these processes. Virtually all neuropsychiatric disorders are associated with sleep dysfunction, including those associated with homeostatic plasticity and Fragile X Syndrome (Bushey et al., 2009; Ferrarelli et al., 2007; Kidd et al., 2014; Sare et al., 2017; Wondolowski and Dickman, 2013; Wulff et al., 2010). Interestingly, sleep behavior is also disrupted by mutations in the *Drosophila* homolog of FMRP, *dfmr1* (Bushey et al., 2009). Further investigation of this provocative network of genes involved in the homeostatic control of sleep and synaptic plasticity may help solve the biological mystery that is sleep and shed light on the etiology of neuropsychiatric diseases.

## MATERIALS AND METHODS

### PHP genetic screen

We identified over 700 mammalian genes that encode transcripts expressed at synapses and that have not been previously screened for PHP. This list was generated from a variety of previous studies and was enriched in putative transcripts associated with FMRP ((Ascano et al., 2012; Brown et al., 2001; Darnell et al., 2011; Gilman et al., 2011; Miyashiro et al., 2003); see Supplementary Table 1)). This list was further supplemented with an additional 176 genes associated with schizophrenia and autism spectrum disorder (Cross-Disorder Group of the Psychiatric Genomics, 2013; Gilman et al., 2011; Jurado et al., 2013; Sando et al., 2012). From these initial lists of mammalian genes, we identified 352 *Drosophila* homologs. We used a combination of known genetic mutations and/or putative transposon mutations (197) or RNA-interference transgenes (341) targeting these genes to obtain a stock collection to screen. Finally, we assessed the lethal phase of homozygous mutants and RNAi lines crossed to motor neuron and muscle Gal4 drivers, removing any mutants that failed to survive to at least the third-instar larval stage. This led to a final list of 134 mutations to screen by PhTx application and 284 RNAi lines to screen by *GluRIII* knock down (Supplementary Table 1). The RNAi screen was performed using T15 and C15 lines, as described (Brusich et al., 2015).

### Fly stocks

All *Drosophila* stocks were raised at 25°C on standard molasses food. The following fly stocks were used in this study: T15 and C15 (Brusich et al., 2015); *inc^1^* and *inc^2^* (Stavropoulos and Young, 2011); *OK371-Gal4* (Mahr and Aberle, 2006); *MHC-Gal4* (Schuster et al., 1996); *UAS-Cul3 RNAi^11861R-2^* ((National Institute of Genetics Fly Stock Center; (Stavropoulos and Young, 2011)); *UAS-dcr2* (Dietzl et al., 2007); *inc-Gal4* (Stavropoulos and Young, 2011); *GluRIIA^SP16^* (Petersen et al., 1997). The *w^1118^* strain was used as the wild-type control unless otherwise noted because this is the genetic background in which all genotypes are bred. *Df*(*1*)*Exel8196* and *UAS-eGFP* were obtained from the Bloomington Drosophila Stock Center. See Supplementary Table 1 for sources of the screened genetic mutants and RNAi lines.

### Molecular biology

*inc^kk^* mutants were generated using a CRISPR/Cas9 genome editing strategy as described (Gratz et al., 2013a; Kikuma et al., 2017). Briefly, we selected a target Cas-9 cleavage site in the first *inc* exon without obvious off-target sequences in the *Drosophila* genome (sgRNA target sequence: 5’ GTTCCTCTCCCGTCTGATTC <u>AGG</u> 3’, PAM underscored). DNA sequences containing this target sequence were synthesized and subcloned into the pU6-BbsI-chiRNA plasmid ((Gratz et al., 2013a); Addgene, Cambridge, MA). To generate the sgRNA, pU6-BbsI-chiRNA was PCR amplified and cloned into the pattB vector (Bischof et al., 2007). This construct was injected and inserted into the attP40 target sequence on the second chromosome (Markstein et al., 2008) and balanced. This line was crossed into a stock expressing Cas9 under control of vas regulatory sequences, which led to 9 independent indels with predicted frameshift mutations in the *inc* open reading frame confirmed by PCR followed by sequencing of *inc* gene in male flies after balancing. Lines which introduced the earliest stop codon (R50Stop) and the second earliest stop codon (C57STOP) were chosen for further analyses and were named *inc^kk3^* and *inc^kk4^* respectively.

To generate *UAS-smFP-inc*, we subcloned the full-length *inc* cDNA from the expressed sequence tag LD43051 (Drosophila Genomics Resources Center; Bloomington, IN) into the pACU2 vector (Han et al., 2011) using standard methods. A spaghetti monster FLAG tag ((10xFLAGsmFP; (Viswanathan et al., 2015)) was PCR amplified and placed in-frame before the stop codon of the *inc* open reading frame. Constructs were sequence verified and injected into the *w^1118^* strain using the VK18 insertion site on the second chromosome (Venken et al., 2006) by BestGene Inc. (Chino Hill, CA). Endogenously tagged *inc^smFP^* was generated by Well Genetics Inc. (Taipei, Taiwan) using CRISPR/Cas9 targetting and homology directed repair (Gratz et al., 2013a; Gratz et al., 2013b). Briefly, a construct containing the smFP-10xFLAG as well as a 3xP3 DsRed reporter was inserted just before the stop codon of the endogenous *inc* locus using a single target gRNA synthesized as RNA and injected. This construct was injected into a *w^1118^* strain with Cas9 expression and the insertion was confirmed by DsRed+ eyes; the DsRed marker was subsequently excised using the pBac transposase, leaving only smFP at the *inc* C-terminus. The insertion was confirmed by genomic PCR sequencing.

### Electrophysiology

Electrophysiology was performed as described (Kiragasi et al., 2017). Electrophysiological sweeps were digitized at 10 kHz with a 1 kHz filter. For all two-electrode voltage clamp (TEVC) recordings, muscles were clamped at −70 mV, with a leak current below 5 nA. mEPSCs were recorded for 1 min from each muscle cell in the absence of stimulation. 20 EPSCs were acquired for each cell under stimulation at 0.5 Hz, using 0.5 msec stimulus duration and with stimulus intensity adjusted with an ISO-Flex Stimulus Isolator (A.M.P.I.). To acutely block postsynaptic receptors, larvae were incubated with or without philanthotoxin-433 (PhTx; 20 μM; Sigma) in HL-3 for 10 mins as described (Dickman and Davis, 2009; Frank et al., 2006). Data were analyzed using Clampfit 10.7 (Molecular Divices), MiniAnalysis (Synaptosoft), Excel (Microsoft), and GraphPad Prism (GraphPad Software).

### Immunocytochemistry

Third-instar larvae were dissected in ice cold 0 Ca^2+^ HL-3 and immunostained as described (Chen et al., 2017). All genotypes were immunostained in the same tube with identical reagents, and then mounted in the same session. The following antibodies were used: mouse anti-Bruchpilot (BRP; nc82; 1:100; Developmental Studies Hybridoma Bank; DSHB); mouse anti-GluRIIA (1:100; 8B4D2; DSHB); guinea pig anti-GluRIID (1:1,000; (Perry et al., 2017)); rabbit anti-DLG (1:5000; (Pielage et al., 2005)); mouse anti-DLG (1:100; 4F3; DSHB); mouse anti-FK1 (1:100; Millipore 04-262); mouse anti-FK2 (1:500; BML-PW8810; Enzo Life Sciences); mouse anti-FLAG (1:500, F1804; Sigma-Aldrich); mouse anti-pCaMKII (1:100; MA1-047; Invitrogen). Donkey anti-mouse, anti-guinea pig, and anti-rabbit Alexa Fluor 488-, DyLight 405-, and Cyanine 3 (Cy3)-conjugated secondary antibodies (Jackson Immunoresearch) were used at 1:400. Alexa Fluor 647 conjugated goat anti-HRP (Jackson ImmunoResearch) was used at 1:200.

### Western blot

Protein extracts were prepared from male whole animals by homogenization in ice-cold NP40 lysis buffer (50mM Tris pH7.6, 150mM NaCl, 0.5% NP40) supplemented with protease inhibitors (Sigma, P8340). Protein lysates were centrifuged at 4°C at 15,000 x g for 15 min and quantitated in duplicate (BioRad, 5000111). 60 μg was resolved by Tris-SDS-PAGE and transferred to nitrocellulose. Membranes were blocked for 1 hr at room temperature in LI-COR Odyssey buffer (LI-COR, 927-40000). Membranes were subsequently incubated overnight at 4 °C in blocking buffer containing 0.1% Tween 20, rat anti-Insomniac (1:1,000) (Stavropoulos and Young, 2011), and mouse anti-tubulin (1:100,000, Genetex, gtx628802). After washing 4×5 min in TBST solution (150mM NaCl, 10mM Tris pH7.6, and 0.1% Tween20), membranes were incubated in the dark for 30 min at room temperature in blocking buffer containing 0.1% Tween 20, 0.01% SDS, Alexa 680 donkey anti-rat (1:30,000, Jackson ImmunoResearch, 712-625-153), and Alexa 790 donkey anti-mouse (1:30,000, Life Tchnologies, A11371). Membranes were washed 4×5 min in TBST, 1×5 min in TBS, and imaged on a LI-COR Odyssey CLx instrument.

### Sleep behavior

One- to four-day-old flies eclosing from cultures entrained in LD cycles (12hr light/12hr dark) were loaded into glass tubes and assayed for 5–7 days at 25°C in LD cycles using DAM2 monitors (Trikinetics). Male flies were assayed on food containing cornmeal, agar, and molasses. Female flies were assayed on food containing 5% sucrose and 2% agar. The first 36–48 hours of data were discarded, to permit acclimation and recovery from CO_2_ anesthesia, and an integral number of days of data (3–5) were analyzed using custom Matlab software (Stavropoulos and Young, 2011). Locomotor data was collected in 1 min bins, and a 5 min period of inactivity was used to define sleep (Huber et al., 2004; Shaw et al., 2000); a given minute was assigned as sleep if the animal was inactive for that minute and the preceding four minutes. Dead animals were excluded from analysis by a combination of automated filtering and visual inspection of locomotor traces.

### Confocal imaging and analysis

Imaging was performed as described (Goel and Dickman, 2018). Briefly, samples were imaged using a Nikon A1R Resonant Scanning Confocal microscope equipped with NIS Elements software using a 100x APO 1.4NA or 60x 1.4NA oil immersion objective. All genotypes were imaged in the same session with identical gain and offset settings for each channel across genotypes. z-stacks were obtained using identical settings for all genotypes, with z-axis spacing between 0.15 μm to 0.5 μm within an experiment and optimized for detection without saturation of the signal. Maximum intensity projections were used for quantitative image analysis with the NIS Elements software General Analysis toolkit. All quantifications were performed for Type Ib boutons on muscle 6/7 and muscle 4 of segments A2 and A3. Type Ib boutons were selected at individual NMJs based on DLG intensity. Measurements were taken from at least ten synapses acquired from at least six different animals. For all images, fluorescence intensities were quantified by applying intensity thresholds to eliminate background signal. For analysis of pCaMKII, BRP, and Inc^smFP^ intensity levels, a mask was created around the DLG channel, used to define the postsynaptic density, and only signals within this mask were quantified. For FK1 and FK2 anti-Ubiquitin staining, mean intensity was calculated using regions within the DLG mask and subtracting intensities from the HRP mask (to exclusively assess the postsynaptic area).

### Statistical Analysis

All data are presented as mean +/− SEM. Data were compared using either a one-way ANOVA and tested for significance using a 2-tailed Bonferroni post-hoc test, or using a Student’s t-test (where specified), analyzed using Graphpad Prism or Microsoft Excel software, and with varying levels of significance assessed as p<0.03 (*), p<0.01 (**), p<0.001 (***), p<0.0001 (****), ns=not significant. For statistical analysis of sleep duration, one-way ANOVA and Tukey-Kramer post hoc tests were used. For all figures, data are quantified as averages +/−SEM, and absolute values and additional statistical details are presented in Supplementary Table 3.

### Data availability

The data that support the findings of this study are available from DD upon reasonable request. The authors declare that the data supporting the findings of this study are available within the paper and its Supplementary Information files.

## AUTHOR CONTRIBUTIONS

KK and DD conceived the project and designed the research. KK, XL, SP, PG, CC, DK, and QL performed experiments. KK, XL, PG, CC, SP, and QL analyzed data. KK and DD wrote the manuscript with feedback from QL, SP, and NS.

## ACKNOWLEDGEMENTS

We thank C. Andrew Frank (University of Iowa, USA), Mark Tanouye (University of California, Berkeley, USA), and Gabrielle Boulianne (University of Toronto, Canada) for sharing fly stocks, and Martin Müller (University of Zürich, Switzerland) for insightful discussions and comments. We acknowledge the Developmental Studies Hybridoma Bank (Iowa, USA) for antibodies used in this study and the Bloomington Drosophila Stock Center for fly stocks (NIH P40OD018537). KK was supported in part by a USC Provost Fellowship. QL was supported by an International Student Research Fellowship from the Howard Hughes Medical Institute (HHMI). This work was supported by a grant from the Mathers Foundation, Whitehall Foundation grant 2013-05-78, fellowships from the Alfred P. Sloan and Leon Levy Foundations, a NARSAD Young Investigator Award from the Brain and Behavior Foundation, the J. Christian Gillin, M.D. Research Award from the Sleep Research Society Foundation, and a Career Scientist Award from the Irma T. Hirschl/Weill-Caulier Trust to NS, and by a grant from the National Institutes of Health (NS091546) and fellowships from the Mallinckrodt, Whitehall, and Klingenstein-Simons Foundations to DD. The authors declare no competing financial interests.

---

**Supplementary Figure 1: *inc* mutants generated by CRISPR/Cas9 gene editing exhibit normal baseline transmission and the expected defects in sleep behavior. (a)** Schematic of the *Drosophila inc* locus. The deleted region of *inc^1^*, the pBac transposon insertion site of *inc^2^*, and the CRISPR-induced early stop codon in *inc^kk3^* and *inc^kk4^* (*) are shown. **(b)** Representative electrophysiological traces of EPSC and mEPSC for wild type (w^1118^), *inc^kk3^, inc^2^, inc^2^/inc^Df^*, and *inc^2^/inc^1^* mutants. While *inc^2^/inc^1^* mutants show reduced synaptic transmission, as reported previously (Li et al., 2017), baseline synaptic transmission is largely normal in the other *inc* alleles. Quantification of average EPSC amplitude **(c)**, mEPSC amplitude **(d)**, mEPSC frequency **(e)**, and quantal content **(f)** values. **(g)** Quantification of average daily sleep in female flies of the indicated genotype. *inc^1^/inc^kk^* females show reduced daily sleep, similar to *inc^1^/inc^2^* transhetrozygotes. **(h)** Expression of UAS-*smFP-inc* driven by *inc-Gal4* restores sleep to wild-type values in *inc^1^* mutants. Error bars indicate ±SEM.

**Supplementary Figure 2: *inc* is required for the chronic expression of PHP. (a)** Representative EPSC and mEPSC traces from the indicated genotypes. PHP fails to be expressed in *inc^kk3^* mutants when combined with loss of *GluRIIA* (*inc^kk3^;GluRIIA^SP16^*). **(b)** Quantification of mEPSC and quantal content values in the indicated genotypes normalized to baseline conditions (*-GluRIIA*).

**Supplementary Figure 3: Overexpression and endogenous tagging of *inc* does not significantly impact baseline neurotransmission or PHP expression. (a)** Representative EPSC and mEPSC traces of wild type, neuronal *inc* overexpression (*OK371-Gal4/UAS-smFP-inc*), and muscle *inc* overexpression (*UAS-smFP-inc/+;MHC-Gal4/+*). Quantification of mEPSC amplitude **(b)**, EPSC amplitude **(c)**, and quantal content **(d)** values from the indicated genotypes. **(e)** Representative EPSC and mEPSC traces from wild type and endogenously tagged *inc^smFP^* before and after PhTx application. Synaptic transmission and PHP function similarly to wild type *inc^smFP^* larvae. **(f)** Quantification of average mEPSC amplitude and quantal content values following PhTx application relative to baseline (-PhTx).

**Supplementary Figure 4: *Cul3* is required postsynaptically for retrograde PHP signaling. (a)** Representative EPSC and mEPSC traces from neuronal *Cul3* knock down (neuronal>*Cul3 RNAi*: *UAS-Cul3 RNAi^11861R-2^/OK371-Gal4*) and muscle *Cul3* knock down (muscle>*Cul3 RNAi*: *UAS-Cul3 RNAi^11861R-2^/+;MHC-Gal4/+*) before and after PhTx application. muscle>*Cul3 RNAi* disrupts the expression of PHP, while PHP expression persists in neuronal>*Cul3 RNAi*. **(b)** Quantification of mEPSC and quantal content values after PhTx application normalized to baseline values. Error bars indicate ±SEM.

**Supplementary Table 1: List of synaptic genes screened and summarized results**. The gene identity (noted by CG number), gene name, putative functions, genotype, genetic perturbation, source, and mEPSP, EPSP, and quantal content values are shown for each mutant screened.

**Supplementary Table 2: Genetic screen identified 17 mutants with reduced basal synaptic transmission**. The CG number, gene name, putative functions, genotype, perturbation type, source, and mEPSP, EPSP, and quantal content values are shown for each mutant identified that exhibited reduced baseline transmission, but robust PHP expression.

**Supplementary Table 3: Absolute values for normalized data and additional details**. The figure and panel, genotype, and conditions used are noted (external Ca^2+^ concentration as well as whether PhTx was applied or not). Average values (with standard error values noted in parentheses) are shown for all data. For electrophysiological experiments, passive membrane properties (input resistance, leak current), mEPSC, EPSC, quantal content (QC), data samples (n), and statistical significance tests and values are shown.

